# Association Analysis and Meta-Analysis of Multi-allelic Variants for Large Scale Sequence Data

**DOI:** 10.1101/197913

**Authors:** Xiaowei Zhan, Sai Chen, Yu Jiang, Mengzhen Liu, William G. Iacono, John K. Hewitt, John E Hokanson, Kenneth Krauter, Markku Laakso, Kevin W. Li, Sharon M Lutz, Matthew McGue, Anita Pandit, Gregory JM Zajac, Michael Boehnke, Goncalo R. Abecasis, Bibo Jiang, Scott I. Vrieze, Dajiang J. Liu

## Abstract

**Motivation:** There is great interest to understand the impact of rare variants in human diseases using large sequence datasets. In deep sequences datasets of >10,000 samples, ∼10% of the variant sites are observed to be multi-allelic. Many of the multi-allelic variants have been shown to be functional and disease relevant. Proper analysis of multi-allelic variants is critical to the success of a sequencing study, but existing methods do not properly handle multi-allelic variants and can produce highly misleading association results.

**Results:** We propose novel methods to encode multi-allelic sites, conduct single variant and gene-level association analyses, and perform meta-analysis for multi-allelic variants. We evaluated these methods through extensive simulations and the study of a large meta-analysis of ∼18,000 samples on the cigarettes-per-day phenotype. We showed that our joint modeling approach provided an unbiased estimate of genetic effects, greatly improved the power of single variant association tests, and enhanced gene-level tests over existing approaches.

**Availability:** Software packages implementing these methods are available at (<u>https://github.com/zhanxw/rvtests http://genome.sph.umich.edu/wiki/RareMETAL</u>).

## 1 Introduction

Rare genetic variants are enriched with functional alleles that play an important role in a variety of complex human diseases, including hematological disorder(Auer, et al., 2014), coronary artery disease(Do, et al., 2015; Myocardial Infarction, et al., 2016; Tg, et al., 2014) and others. The discovery of such rare variant associations has contributed significantly to the generation of new mechanistic insights and the identification of novel therapeutic targets(Cohen, et al., 2006; Tg, et al., 2014). These discoveries are critical steps toward the successful implementation of precision medicine.

As the cost of sequencing continues to decrease, many sequence-based studies of rare variants have begun to emerge. Compared to array-based studies which only genotype variants at known sites, sequence-based studies unbiasedly reveal both known and novel variants across the frequency spectrum. The fraction of novel alleles/variants uncovered increases with increasing read depth and sample size. In addition to identifying novel variant sites, numerous novel alleles at known variant sites are being uncovered as well. As shown in the exome aggregation consortium (ExAC)(Lek, et al., 2016), 8% of the variant sites in the human exome are multi-allelic and contain more than one alternative allele. A number of these multi-allelic variants are functional and have been shown to be disease relevant (Lek, et al., 2016). Despite the importance of multi-allelic variants, most of the methods developed so far for sequence-based association analysis consider only bi-allelic variants, and thus do not properly handle multi-allelic sites (Chang, et al., 2015; Purcell, et al., 2007).

The analysis of multi-allelic sites is currently often ignored in GWAS and sequence-based association studies. Multi-allelic analysis were considered prior to GWAS era for microsatellite markers. Yet, the existing methods all have certain limitations, which make it challenging to analyze sequence data. Some methods focused on how to combine multiple alleles in the same position and perform an omnibus test(El Galta, et al., 2005). Another method (Terwilliger, 1995) made use of retrospective likelihood to model the joint distribution of multi-allelic variants at a single variant site. Yet it is challenging to extend this model to multiple variant sites in linkage disequilibrium, it is difficult to generalize this approach to analyze gene-level associations. To our knowledge, it is still unknown what the best strategy is to integrate multiple allelic sites into gene-level association tests.

Moreover, it is unclear how to perform meta-analysis and combine samples across studies in the presence of multi-allelic variants. In addition, most of the identified rare variant associations have small to moderate effect sizes(Zuk, et al., 2014). There is growing recognition that large sample sizes are needed to attain sufficient power to uncover rare causal variants. Consortia efforts are underway to aggregate large sample sizes for the study of various complex human diseases. Meta-analysis plays a critical role in the vast majority of consortium efforts, where typically only summary level information such as genetic effects and p-values are shared across different studies. Compared to sharing individual-level genotype and phenotype data from study participants, meta-analysis of summary statistics can be easier to implement, more protective of study participant privacy and more robust against heterogeneity between studies(Evangelou and Ioannidis, 2013). It is therefore necessary to extend existing meta-analysis methods and software to properly handle multi-allelic sites as well.

In this article, we propose a series of innovations to address the key analysis issues for multi-allelic variants, which represent 10% for the genomic variations. We developed novel methods to jointly model the effects of multiple alleles in single variant association tests, and facilitate convenient gene-level association analysis and meta-analysis. We evaluated these methods using extensive and realistic simulations and show that they consistently outperform existing naive methods that either ignore multi-allelic sites or test each alternative allele separately. We also applied these methods to a large scale meta-analysis of nicotine addiction phenotypes. We show that our method can uncover multi-allelic association in known loci of the cigarettes-per-day (CPD) phenotype. We have also implemented these methods in RVTESTS(Zhan, et al., 2016) for association analysis and the generation of summary association statistics and RAREMETAL(Feng, et al., 2014) for meta-analysis. Given the importance of the multi-allelic variants, we expect these methods to play key roles in large scale genetic discoveries with sequence data.

## 2 Methods

We describe our method to encode multi-allelic variants, perform single variant and gene-level analyses, and carry out meta-analysis. The key idea is to jointly model the effects of multiple alternative alleles for multi-allelic variants in single variant and gene-level tests. This joint modeling strategy gives a proper estimate of the alternative allele effect and facilitates the construction of gene-level tests from single variant association test statistics of multi-allelic sites. This method improves power over the method that ignores multi-allelic variants and the method that models the effect of each alternative allele separately.

For a multi-allelic variant at site *m* with *L* alternative alleles, we can encode the genotype for individual *i* with an *L*-vector 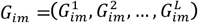, where the *l*^*th*^ entry is the number of the *l*^*th*^ alternative allele. Assuming Hardy-Weinberg equilibrium, the counts 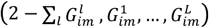 follow a multinomial distribution *Multinom* (2, ((1 —Σ_*l*_ *f*_*1*_),*f*1,…,*f*_*L*_)), where *f*_*1*_ is the alternative allele frequency for the *I*^*th*^ alternative allele, and 1 — Σ_*l*_ *f*_*1*_ is the frequency for the reference allele. The counts for two different alternative alleles 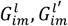 are negatively correlated with covariance 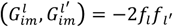.The correlation can be large when the two alternative alleles *A*_*l*_ and 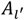 are common. We have illustrated this genotype coding with an example of a tri-allelic site in **Table S1.**

When there are genotype uncertainties in the data, genotype dosages are often used instead of hard genotype calls for genetic association analyses(Howie, et al., 2012; Howie, et al., 2009; Li, et al., 2011). Under our coding scheme, the calculation of the genotype dosages is similar to biallelic variants.

### Joint Modeling Multi-Allelic Effects

We are interested in estimating and testing for the effect of each alternative allele *A*_*l,*_ *l =* 1, *L.* The effect of allele *A*_*l*_ measures the mean phenotype change when having an additional copy of the *A*_*l*_ allele.

To properly analyze a multi-allelic variant, we propose a joint model that includes the genotypes for all alternative alleles in the model. Specifically, to estimate (or test for) the effect of the *l*^*th*^ alternative allele, we perform the multiple regression *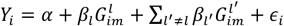.* The multiple regression coefficient β_*l*_ estimates 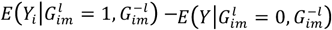 where 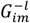 is the genotype vector at site m for the rest of the alleles *A*_1_,…, *A*_*l*-1_, *A* _*l*+1_,…, *A*_L_. The effect of the *l*^*th*^ alternative allele can be unbiasedly estimated from multiple regression.

An alternative strategy, which we call single-allelic analysis, is to restrict our analysis to the set of individuals with genotypes *A*_*0*_*/A*_*0*_ *,A*_*0*_*/A*_*l*_ *,A*_*l*_*/A*_*V*_ As the analyzed samples are selected based on genotype only, the regression analysis is still valid and will give us an unbiased estimate of the effect of *A*_*t*_. However, depending on the frequency of other alternative alleles, the single-allelic analysis may discard a significant portion of the sample and the association analysis can be underpowered.

An additional advantage of joint multi-allelic analysis over single-allelic analysis is the convenience of constructing gene-level tests from single variant association statistics. For single-allelic analysis, a different set of samples are analyzed for each different alternative allele. This makes it impossible to construct gene-level tests using single variant association statistics calculated for different samples.

Finally, it is important to note that directly regressing *Y* over the allele count 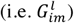 will lead to biased effect estimates. A numerical example is given in the **Supplemental Methods** and **Figure S1,** to illustrate the considerable bias and inflated type I errors for a naïve approach.

### Meta-analvsis of Single Variant Test in the Presence of Multi-allelic Sites

We propose appropriate meta-analysis methods of single variant and gene-level association tests in the presence of multi-allelic sites. We denote the sample genotype matrix at the multi-allelic site *m* as **G_m_.** We will calculate and share the marginal association statistic obtained from the regression analysis over the counts of each alternative allele, i.e. 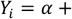 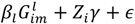, where *Z*_*t*_ is the vector of covariate for individual *i.* The score statistic for the *l*^*th*^ allele is equal to 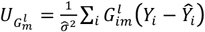 where *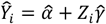.* The parameters 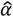 and 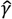 are the model parameter estimates for *α* and *γ* and 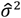 is the residual variance under the null hypothesis. The variance-covariance matrix between the score statistics for different alleles are given by

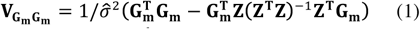

To test for the effect of the *l*^*th*^ alternative allele, we need to control for the effects of the rest of the *L —* 1 alternative alleles. Specifically, in a regression model that includes the counts of all alternative alleles, i.e.

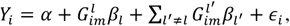

the conditional score statistic for the *I*^*th*^ alternative allele is equal to 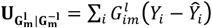, where 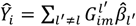 and 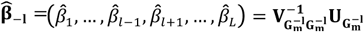

The conditional score statistic can be calculated using marginal association statistics:

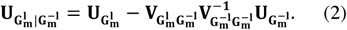

The variance of the conditional score statistic is equal to

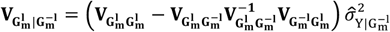

In meta-analysis, we will combine score statistics using the Mantel Haenszel method. Specifically, given the score statistics at site *m* 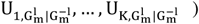 and their variances in *K* studies 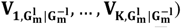, the meta-analysis score statistic can be calculated by 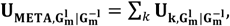 and 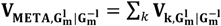.

The standardized score statistic is equal to 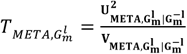, which follows a chi-square distribution with 1 degree of freedom.

### Meta-analvsis of Gene-level Association Test in the Presence of Multi-allelic Sites

As we showed for single variant analysis, it is necessary to jointly model the effects of all alternative alleles in the same site in order to attain unbiased association analysis of each allele. Most commonly used gene-level association tests, such as the burden test, SKAT and VT, can be constructed using single variant association statistics and their covariance matrices(Lee, et al., 2013). When the gene region contains rare alternative alleles from multi-allelic sites, the score statistic from joint multi-allelic analysis 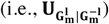 needs to be used to construct a gene-level test. As in single variant analysis, using the marginal score statistic 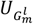 without adjusting the effects of other alternative alleles leads to biased results and inflated type I errors.

Below, we describe an extension of gene-level tests to scenarios where the gene region contains multi-allelic sites. The calculation of gene-level tests requires score statistics from variant sites that contain rare alleles, including the score statistics from bi-allelic sites, the score statistic from joint multi-allelic analysis, as well as the covariance matrix between them. Single variant association statistics from bi-allelic and multi-allelic sites have been described in the above section. We next derive the variance-covariance matrix between these score statistics and then discuss how to use them to construct commonly used rare variant tests.

For notational convenience, we denote the genotype matrices for common alternative alleles from multi-allelic sites as **G_c_,** the rare allele from the multi-allelic sites as **G_R_** and the rare alleles from bi-allelic sites as **G_b_.** We denote the vector of score statistics for all rare alleles as 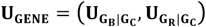, which includes the score statistics from bi-allelic sites (conditional on the common alternative alleles from multi-allelic sites) and the score statistics from joint multi-allelic analysis.

Below, we illustrate how to calculate the covariance matrix between score statistics. The covariance matrix between score statistics of rare alleles at multi-allelic sites equals to

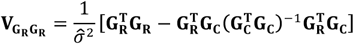

the covariance between rare bi-allelic variants equals to

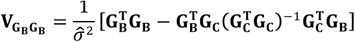

The covariance matrix between rare bi-allelic variants and rare multi-allelic variants equals to

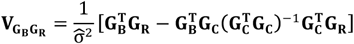

When non-genetic covariates **Z** are present, we just need to replace **G_c_** with 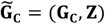, and the calculation of covariance matrix remains the same.

The covariance matrix for the score statistic is denoted by 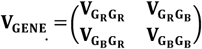 The burden test statistic(Li and Leal, 2008) and its variance are equal to 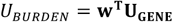 and 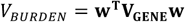, where w is the weight assigned to each variant. The standardized burden statistic 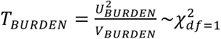. The SKAT statistic(Wu, et al., 2011) is equal to 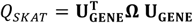, where Ω is a diagonal matrix with the diagonal entries being the weights assigned to each variant site. The SKAT statistic follows a mixture chi-square distribution with mixture proportions being the eigenvalues for 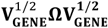. The VT statistic calculates a burden test statistic for each minor allele frequency threshold and corrects for the multiple comparison issue using the minimal p-value method(Lin and Tang, 2011; Price, et al., 2010). The p-values can be calculated using the distribution function for a multivariate normal distribution.

### Design of Simulation Evaluation

We conducted extensive simulations to evaluate the proposed methods. To generate genetic data with realistic patterns of multi-allelic sites, we used the allele frequency spectra estimated from large-scale exome sequencing projects. We downloaded data from the ExAC project (version 0.3.1), which consists of summary information for coding variants from 60,706 exomes.

To benchmark single variant association tests, in each replicate, we randomly picked one variant site from 219,680 sites that contain multiple alternative alleles. To illustrate the advantage of joint multi-allelic analysis, we separately considered the power for detecting associations with the primary alternative allele (i.e. the most frequent alternative allele) and the secondary alternative allele (i.e. the less frequency allele(s)). We simulated the genotype (i.e. the reference and alternative allele counts) for each sample based on a multinomial distribution: *multinom* (2, ((1 —Σ_*l*_ *f*_*1*_),*f*1,…,*f*_*L*_)). For each variant site, we randomly chose one alternative allele as causal with effects being 0.1, 0.25 or 0.5 sd. The power for detecting associations with the primary (or secondary) alternative allele was assessed by the fraction of the replicates with significant p-values (<5xl0∼ ^8^) among the replicates where the primary (or secondary) alternative allele is causal.

We also assessed the power for single allelic and joint-multi-allelic analysis as omnibus tests to identify associated variant sites (instead of identifying associated alleles). We compared it with the off-the-shelf method of collapsing the multiple alternative alleles.

In order to evaluate the gene-level association test under the most realistic patterns of linkage disequilibrium and multi-allelic variant allele frequency spectrums, we made use of real genotype data from eight cohorts included the Minnesota Center for Twin and Family Research (MCTFR), SardiNIA, METabolic Syndrome In Men (METSIM), Genes for Good, COPDGene with samples of European ancestry, COPDGene with samples of African American ancestry, and the Center for Antisocial Drug Dependence (CADD). Based upon the real genotype, we simulated phenotypes: for each replicate, we randomly chose one gene with at least one multi-allelic variant. We chose a fraction (20% or 50%) of the genetic variants as causal, with effects simulated from iV(0,0.2^2^). We considered three commonly used tests, including the simple burden test, SKAT and VT. The type I error and power for the meta-analyses were evaluated under *α=* 2.5xl0^-6^.

### Analysis of Cigarettes Per Day Phenotype (CPD)

In order to benchmark our method and its implementation, we applied our method to perform a meta-analysis on large genetic datasets from eight cohorts for the CPD phenotype. The eight cohorts included the Minnesota Center for Twin and Family Research (MCTFR), SardiNIA, METabolic Syndrome In Men (METSIM), Genes for Good, COPDGene with samples of European ancestry, COPDGene with samples of African American ancestry, and the Center for Antisocial Drug Dependence (CADD). Summary association statistics from the eight cohorts were generated using RVTESTS(Zhan, et al., 2016), and meta-analysis was performed centrally using RAREMETAL(Feng, et al., 2014). Detailed descriptions of the cohorts are available in **Supplemental Methods Section 2,** including information on the methods for association analyses and the adjusted covariates.

In order to ensure the validity of our association analysis results, we conducted extensive quality control for the imputed genotype data. We filtered out variant sites with the imputation quality metric *R*^2^ < .7, and removed variant sites that showed large differences in allele frequencies from the reference panel. We performed single variant tests using joint multi-allelic analysis and single-allelic analysis. We also performed genelevel tests using the burden test, SKAT and VT under two different allele frequency cutoffs, 1% and 5%. As a comparison, we analyzed the data using the method that discards the multi-allelic sites as well.

## 3 Results

### Type I Error and Power Evaluation for Single Variant Association Test

Simulations indicated that jointly modeling the allelic effects of multiple alternative alleles leads to more powerful single variant association tests (Table 1). The power for joint multi-allelic analysis is consistently higher than single allelic analysis. We separately considered the power for the analysis of the primary and secondary alternative alleles. For the analysis of secondary alternative alleles, the single allelic analysis did not consider samples that carry the primary alleles. The power for single allelic analysis was much lower than multi-allelic analysis. For example, in the scenario where the causal allele effect is .25, the power for single allelic analysis is .36 whereas the power for multi-allelic analysis is .43. On the other hand, the power of multi-allelic analysis for detecting associations with primary alternative allele has a smaller advantage.

**Table 1.** The power for single variant association analysis. We compared the power of single allelic analysis and joint multi-allelic analysis for detecting associations with each alternative allele. The power was evaluated under the threshold of *a* = 4.5xl0^-8^, adjusting for the increased multiple testing burden for analyzing multiple alleles.

| Sample Size | Genetic Effects | Single Allelic Analysis | Multi-Allelic Analysis |
| --- | --- | --- | --- |
| **Type I Error/Power for the Analysis of the Primary Alt Alleles** |  |  |  |
| 10000 | 0 | $4.7 \times 10^{-8}$ | $4.2 \times 10^{-8}$ |
|  | 0.1 | 0.24 | 0.25 |
|  | 0.25 | 0.57 | 0.57 |
|  | 0.5 | 0.75 | 0.76 |
| 20000 | 0 | $4.6 \times 10^{-8}$ | $4.2 \times 10^{-8}$ |
|  | 0.1 | 0.36 | 0.37 |
|  | 0.25 | 0.67 | 0.68 |
|  | 0.5 | 0.82 | 0.82 |
| **Type I Error/Power for the Analysis of Secondary Alt Alleles** |  |  |  |
| 10000 | 0 | $4.1 \times 10^{-8}$ | $4.8 \times 10^{-8}$ |
|  | 0.1 | 0.037 | 0.056 |
|  | 0.25 | 0.24 | 0.3 |
|  | 0.5 | 0.48 | 0.55 |
| 20000 | 0 | $4.9 \times 10^{-8}$ | $4.3 \times 10^{-8}$ |
|  | 0.1 | 0.087 | 0.12 |
|  | 0.25 | 0.36 | 0.43 |
|  | 0.5 | 0.6 | 0.66 |

We also compared the power for the single allelic and multi-allelic analysis as omnibus tests for identifying associated variant sites **(Table S2).** Testing each allele separately may slightly increase the burden for multiple testing. In a deep sequencing study, 10% of the variant site can be multi-allelic. Using single allelic or multi-allelic analysis as omnibus test, a variant site is deemed to be associated if at least one alternative allele has p-values < 5xl0^-8^/l.l, a threshold that corrects for the increased load of multiple testing. The power for the collapsing method was evaluated under the threshold of 5x10^-8^. We considered models where 1) only the primary alternative allele is causal, 2) only secondary alternative is causal, and 3) the model where all alternative alleles are causal. The power for single allelic and multi-allelic analysis is higher than the method that collapses multiple alleles under nearly all scenarios. When all alternative alleles are causal with effects in the same direction, the collapsing method is only slightly more powerful. When only the secondary alternative is causal, the presence of non-causal primary alternative allele can severely weaken the association signal and substantially reduce the power for the collapsing method.

### Type I Error and Power Evaluation for Gene-level Association Test

We evaluated the power for two different analysis strategies for gene-level tests in the presence of multi-allelic sites: 1) the joint modeling approach that simultaneously considers the effects of multi-allelic and bi-allelic sites, and 2) the approach that discards multi-allelic sites from the gene-level analysis.

We also evaluated the power under a variety of scenarios with different combinations of sample sizes, genetic effect distributions and proportions of causal variants. Causal variant effects were sampled from a normal distribution 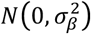, with 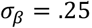. Under each scenario, three gene-level tests were considered: the simple burden test, SKAT, and VT, analyzing rare variants with MAF<1% or *5%.*

Type I errors were well controlled across all scenarios. The power for gene-level tests was consistently higher when we jointly modeled the effects of all alternative alleles for multi-allelic sites (Table 2). The strategy that discards multi-allelic sites could lead to ∼20% decrease in power, particularly when the effects of causal alleles are in the same directions. For example, when the MAF cutoff of 0.01 and 20% of the variants were causal, power for the burden/SKAT/VT tests were respectively 50%/39%/68%, which were substantially higher than the power for the three tests analyzing only bi-allelic variants (42%/35%/61%). This is consistent with the benchmark of rare variant association methods (Li and Leal, 2008; Liu and Leal, 2010), where the erroneous exclusion of causal variants drastically reduces power.

**Table 2.**
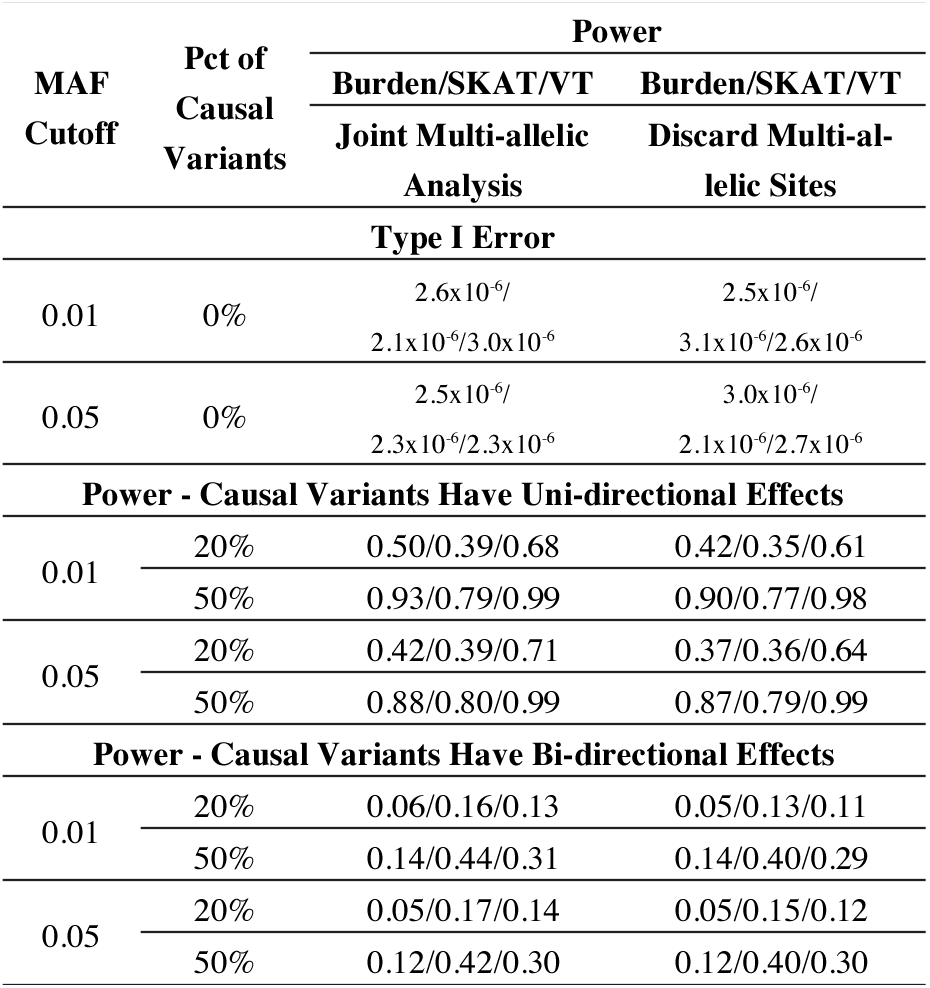
The Type I Error and Power for Gene-level Association Tests. We compared the power for simple burden, SKAT and VT tests for the joint multi-allelic analysis and the analysis that discards multi-allelic sites. The power and type I error were assessed under a threshold of *a =* 2.5xl0^-6^ using 100,0000 replicates.

When variants have bi-directional effects, SKAT is the most powerful test. The power of SKAT based upon multi-allelic analysis was considerably higher than the method that discards multi-allelic sites. Simple burden and VT tests were underpowered in this scenario. Yet, the tests based upon multi-allelic analysis were still consistently more powerful.

### Analysis of Cigarettes-Per-Day Phenotype

We analyzed the genetic and phenotype data from the eight cohorts for the study of the CPD trait. The eight cohorts were genotyped with GWAS arrays, and imputed to the 1000Genome reference panel with the Michigan Imputation Server(Das, et al., 2016; McCarthy, et al., 2016). After quality control, a total of 29,124,949 variants sites were segregating in at least one cohort. Among them, 289,809 (1%) contained multiple alternative alleles. The fraction of multi-allelic sites was lower than what was discovered in sequence-based studies, due to the sample size of the reference panel, the exclusion of the variant sites with low alternative allele counts from the reference panel, and the removal of rare imputed alleles due to low imputation quality. Most of the multi-allelic variants were in the intergenic region, and only 105,727 belonged to the genic region. Among the 19,321 genes that were analyzed, 2,475 contained coding multi-allelic variants (nonsynonymous, stop or splice), and 2,319/2,417 contained rare alternative alleles with MAF<1%/5% at their multi-allelic sites.

We first performed single variant association tests. The analysis results are well behaved. We examined the genomic control values for all variants in different frequency bins (0, 0.001], (0.001, 0.01] and (0.01, 0.5], All genomic control inflation factors were < 1.03. We also separately examined the genomic control inflation factor for multi-allelic variants only, and ensured that the tests all had well-calibrated type I errors **(Figure S3).**

In single variant association analysis, we recovered a well-known locus associated with CPD. In fact, all top hits in the meta-analysis came from the *CHRNA5-CHRNA3-CHRNB4* locus. The top variant was 15:78886947_G/A (rs4887067), which is a variant from the untranslated region in the gene *CHRNA5.* No other loci, or novel loci, were uncovered in this study with genome-wide significance.

The top association signals for multi-allelic variants also lay in the *CHRNA5-CHRNA3-CHRNB4* locus (Table 3, **Figure S2-S4).** Most of the top association signals appeared as insertion-deletion polymorphisms (indels). The most significant variant 15:78915370(rs34573245) had a p-value of 1.6x10 ^11^ and is located in the intergenic regions. There were five other significantly associated multi-allelic variants in the genes *CHRNA5, CHRNA3* and *IREB2.*

**Table 3.**
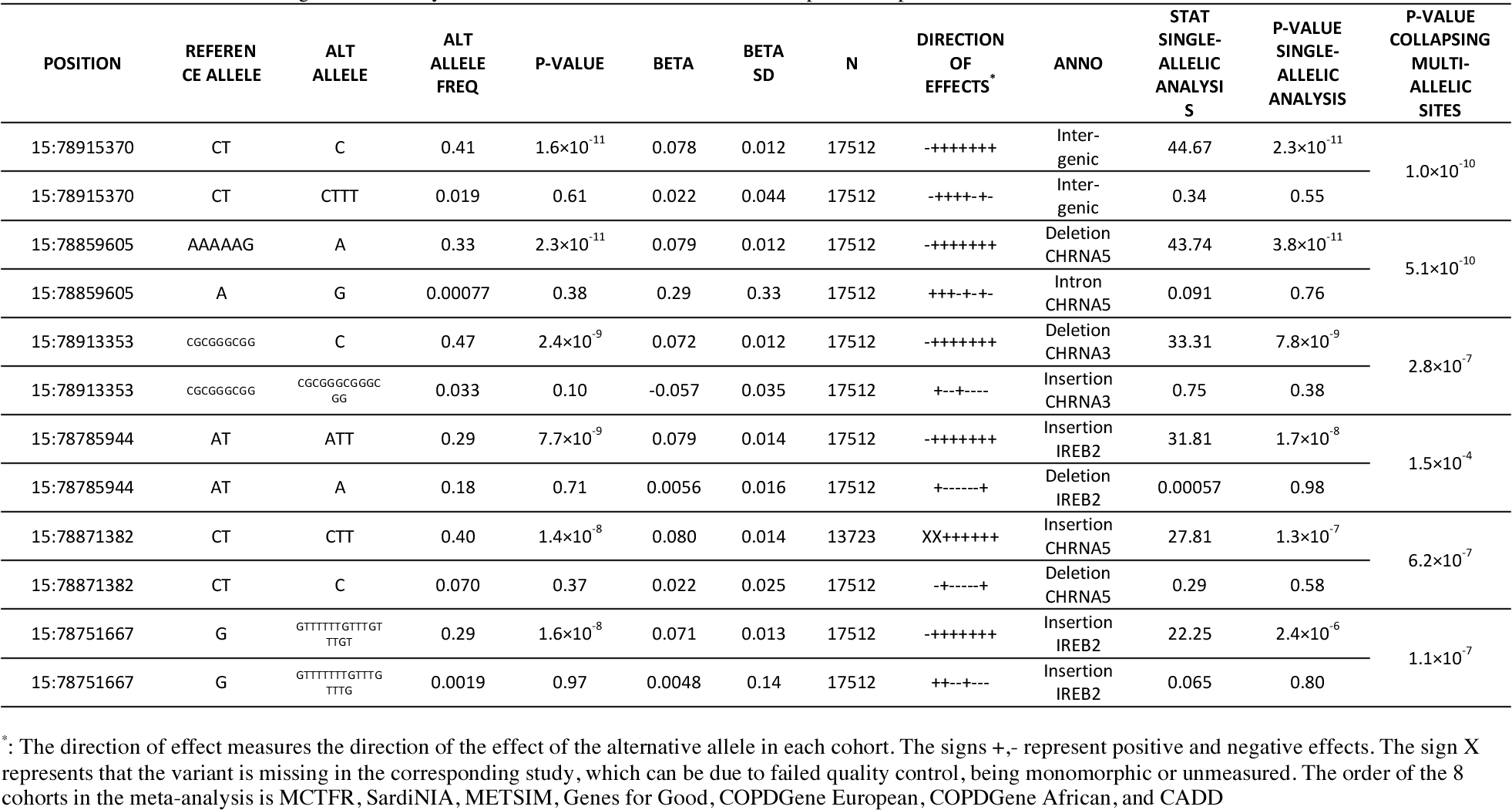
Top Single Variant Association Signals for the Cigarettes-Per-Day Phenotype Using Multi-allelic Analysis. Results are shown for variants with p-values less than 5x10^8^. We report the p-values and genetic effect estimates for each alternative allele at multi-allelic sites. As a comparison, we also report the p-values and test statistics from single-allelic analysis, as well as the omnibus test that collapses multiple alleles.

We compared the association results using the new method and the method that relies on single-allelic analysis. Single-allelic analysis identified only 5 significantly associated multi-allelic variants in the locus, while the joint multi-allelic analysis identified 6 variants. The p-values for multi-allelic analysis were consistently smaller. The mean chi-square statistic at known loci could be used as an estimate for the non-centrality parameter and then used as a metric to empirically assess the power for an association statistic(Zaitlen, et al., 2012). The mean chi-square statistics for the multi-allelic and single allelic analyses were 34.5 and 33.3 respectively, with multi-allelic variants 4% higher. This is consistent with the observations in our simulation studies. We plotted the — log_10_(P) for the two methods **(Figure S1).** We observed a higher concordance between multi-allelic and single-allelic association analysis for lower frequency variants (MAF<1%) than for common variants (MAF>1%), with rank correlations for common and rare variants at 98% and 90% respectively.

In addition, we also implemented the method that collapses multiple alternative alleles. The collapsing method is an omnibus test, which can be used to identify associated variant sites, instead of associated alleles. Given that all the top association signals are driven by the common primary alternative allele, all collapsing p-values were less significant than multi-allelic analysis p-values. No additional significant variant sites were identified. Among the 6 top variant sites identified using joint multi-allelic analysis, only two variant sites 15:78915370 and 15:78859605 had ge-nome-wide significant collapsing p-values (p < 5xl0^-8^).

We also performed gene-level association tests analyzing variants with MAF<1% and 5%. Type I errors were well controlled for all gene-level tests **(Figure S5, S6).** For genes with rare multi-allelic variants, no significant associations were found (Table 4). Only one gene *SHCBP1L* was identified as significant under the Bonferroni threshold a=2.5xl0^6^ for testing 20,000 genes **(Table S3).** The gene is a testis-specific spindle-associated factor that plays a role in spermatogenesis(Liu, et al., 2014; Sood, et al., 2001), which does not have an obvious function related to tobacco use phenotypes.

**Table 4.** Top Gene-level Association Signals for Genes with Multi-allelic Sites. We performed simple burden, SKAT and VT tests under the two different minor allele frequency cutoffs 0.01 and 0.05. No results were significant under the threshold *a =* **2.5x 10^-6^**. For each rare variant test performed, we show the test statistics, p-values, the number of variant sites and the number of multi-allelic variant sites for the top 3 signals.

| Gene | Statistic | P-Value | Number of Variant Site | Number of Multi-allelic Site | Number of Multi-allelic Site with Rare Variant | Gene | Statistic | P-Value | Number of Variant Site | Number of Multi-allelic Site | Number of Multi-allelic Site with Rare Variant |
| --- | --- | --- | --- | --- | --- | --- | --- | --- | --- | --- | --- |
| Burden Test with MAF<1% |  |  |  |  |  | Burden Test with MAF<5% |  |  |  |  |  |
| MLKL | 16.81 | $4.1 \times 10^{-5}$ | 28 | 4 | 4 | PTPN22 | 13.49 | 0.00024 | 32 | 3 | 1 |
| DMBX1 | 13.11 | 0.00029 | 15 | 2 | 2 | CROCC | 11.14 | 0.00085 | 178 | 1 | 1 |
| BRD3 | 10.73 | 0.0011 | 29 | 3 | 2 | HLA-DQA1 | 10.25 | 0.0014 | 11 | 3 | 0 |
| SKAT Test with MAF<1% |  |  |  |  |  | SKAT Test with MAF<5% |  |  |  |  |  |
| ABTB1 | 1654137.32 | $5.5 \times 10^{-5}$ | 28 | 3 | 3 | ABTB1 | 1834395.76 | 0.00015 | 29 | 3 | 3 |
| SEMA7A | 1056004.78 | 0.00032 | 13 | 1 | 1 | DTNBP1 | 4098263.16 | 0.00049 | 23 | 1 | 1 |
| METTL8 | 1075541.99 | 0.00036 | 10 | 9 | 8 | NRBF2 | 1454883.06 | 0.00074 | 7 | 4 | 4 |
| VT Test with MAF<1% |  |  |  |  |  | VT Test with MAF<5% |  |  |  |  |  |
| TTC15 | 21.98 | $1.9 \times 10^{-5}$ | 27 | 18 | 13 | TTC15 | 21.98 | $2.46 \times 10^{-5}$ | 27 | 18 | 15 |
| MLKL | 16.81 | 0.00031 | 28 | 4 | 4 | WNK1 | 16.11 | 0.00049 | 28 | 15 | 15 |
| ARHGEF40 | 15.08 | 0.00078 | 28 | 3 | 1 | MLKL | 15.71 | 0.00068 | 28 | 4 | 4 |

Finally, we compared the gene-level test p-values for the analysis that discards multi-allelic sites, and the analysis that only analyzes rare alternative alleles without controlling for the genotypes of the common alternative allele in the same site. Considerable discrepancies were observed in the scatterplots for different analysis strategies **(Figure S7),** which shows that the naïve method does not provide a useful approximation for the principled methods.

Our software implementation scaled well with this large-scale analysis. The generation of single variant association statistics took 15.1 CPU hours. The computing time scales linearly with the sample size and the number of genetic variants. It required 2.1 CPU hours for single variant meta-analysis and 6.2 CPU hours for three gene-level association tests conducted under two different MAF cutoffs.

## 4. DISCUSSION

Multi-allelic variants represent a highly important class of genetic variation in large scale sequencing studies. Multi-allelic variants have been largely ignored in the GWAS era due to the extensive use of common bi-allelic SNPs as markers to tag regions that harbor causal variants. As deep sequence data become increasingly available on larger sample sizes, many more low frequency and rare multi-allelic variants are expected to be discovered. Many of the novel variants will be identified at previously monomorphic sites, while others will appear as novel alleles at known sites. For example, in a sample with ∼65,000 exomes, 8% of the known variant sites were found to be multi-allelic. It will be important to be able to properly analyze such multi-allelic variants for disease association and assess their functional impact.

Here, we developed and evaluated a new method for analyzing multi-allelic sites in sequence-based association studies and meta-analysis. The method proceeds by jointly modeling the effects of different alternative alleles for multi-allelic variants. It allows unbiased estimates of multi-allelic effects and leads to more powerful gene-level tests than the method that discards multi-allelic variants.

Most of the multi-allelic variants from imputation-based GWAS contain no more than 2 different alternative alleles. For single nucleotide variants, there can be at most 3 different alternative alleles. We focus on testing the effect of each alternative allele for association, while jointly modeling the effects of other alternative alleles. This strategy appears to be the most biologically relevant. We are interested in knowing if a given basepair change is associated with the phenotype, so that we can follow up with precise functional experiment to validate these discoveries.

For the analysis of other types of variants such as indels or copy number variations, there may be more alternative alleles. It may be of interest to perform an omnibus test (e.g. by collapsing multiple alternative alleles) and examine if at least one allele at the site is associated with the disease outcome. One possibility is to consider multivariate tests, such as the multivariate score test(Hotelling, 1931), the method by collapsing multiple alternative alleles, or the variance component score based test (Lin, 1997). It is well known that these multivariate tests can be calculated using the shared summary association statistics and their covariance matrices. Thus, our framework can be easily adapted to omnibus tests for multi-allelic variants. It should be noted that the omnibus test can be extremely underpowered if only the secondary alternative alleles are causal. As most of the novel alternative allele identified by a sequencing study is rare, the utility of omnibus test in the association of multi-allelic variants remain to be understood in the upcoming large scale sequencing studies.

As an application, we applied our method to a large scale meta-analysis of the cigarettes-per-day phenotype using the 1000 Genomes Project based imputations. The analysis type I error rates were well-behaved and sufficiently powerful, confirming the known association for the *CHRNA5-CHRNA3-CHRNB4* locus. Yet, no new loci were uncovered from our meta-analysis using ∼18,000 samples. This may be because our dataset is smaller than some of the largest studies on tobacco addiction(Tobacco and Genetics, 2010). It is clear that larger sample sizes may be necessary to uncover novel nicotine addiction-related loci.

As multi-allelic variants are often ignored by existing GWAS and sequence-based association analysis software packages, the representation of multi-allelic variants is still not unified. The output from the popular imputation software and imputation servers(Das, et al., 2016) represent multiple alleles from the same site in separate lines, with the genotypes in each line representing the number of the corresponding alternative alleles. For instance, the variant at chromosome 19 and position 55178198 has reference allele C, and two possible alternative alleles G and T. The VCF file contains two lines for this variant site: one line with reference/alternative alleles being C/G and the other line with reference/alternative alleles C/T. To represent individual genotypes, one must combine information across the two lines in the VCF. For example, genotypes of G/T (i.e., heterozygous for both alternative alleles) would be represented with a genotype coding of 0/1, 0/1 in the two lines of VCF file. Similarly genotypes of C/G would be encoded as 1/0 and 0/0. In other VCFs, such as the VCF files released by the ExAC project(Lek, et al., 2016), the multi-allelic variant may be represented in one single line. For instance, the same variant 19:55178198, 0,1 and 2 may be used to represent the reference allele C, the first and second alternative alleles G and T. In this case, the genotype G/T is coded as 1/2. Software packages are available to recode the genotypes of multi-allelic site in separate lines. Our implementation of the method supports both representations. In the future, it will be helpful to standardize the representation of the multi-allelic sites and streamline the support in software packages and libraries.

In conclusion, we developed a series of methods for multi-allelic association analysis and meta-analysis, which provide unbiased effect estimates for multi-allelic variants and improve power over current available approaches. As large scale sequencing studies become more prevalent, multi-allelic variants will become an even more important class of genetic variation. We envision that our methods will be highly applicable for understanding the functional impact and disease associations of multi-allelic variants in large scale sequencing studies.

